# Yeast surface display reveals structure-dependent β-glucan association with the plant immune receptor *At*IGP1

**DOI:** 10.64898/2026.10.28.762498

**Authors:** Benjamin Menn, Antonio Molina, Lucía Jordá Miró, Markus Pauly, Vicente Ramírez

## Abstract

Cell wall-derived oligosaccharides can function as immune-active signals in plants and trigger pattern-triggered immunity (PTI) through pattern recognition receptors (PRRs). In *Arabidopsis thaliana*, the leucine-rich repeat malectin (LRR-MAL) receptor kinase IGP1 (impaired in glycan perception 1) contributes to immune responses induced by cellulose- and mixed-linkage β-1,3/1,4-glucan (MLG)-derived oligosaccharides, although whether MLG-derived oligosaccharides are directly recognized by *At*IGP1 needs further clarification. We established a *Saccharomyces cerevisiae* surface-display system in which the *At*IGP1 extracellular domain (ECD) is presented at the yeast cell surface, allowing oligosaccharide recognition to be examined separately from downstream plant signaling and without receptor purification. Previously, cellulose-derived oligosaccharides have been shown to elicit plant defense responses via *At*IGP1. Here, surface-displayed *At*IGP1 retained the ability to recognize fluorescently labeled cellotriose (CEL3) and cellopentaose (CEL5) ligands, supporting functional presentation of the receptor for investigating glycan recognition in a heterologous context. Using this platform, we investigated the association of *At*IGP1 with fluorescently labeled MLG-derived oligosaccharides differing in degree of polymerization (DP) and glycosidic linkage arrangement. This analysis revealed a specific, competitively displaceable association with MLG DP2 (G3G; laminaribiose), whereas the DP3 (G4G3G) and DP4 (G4G4G3G) structures tested showed no detectable association. To test whether this association is relevant *in planta*, we treated *A. thaliana* seedlings with unlabeled MLG DP2, which induced a cytosolic Ca²⁺ response, an early readout of immune activation. This response was markedly reduced in the *igp1-3* mutant. Together, these results identify laminaribiose as an additional candidate ligand of *At*IGP1. The established yeast surface-display platform thus provides an approach for testing candidate oligosaccharides for association with the extracellular domains of plant PRRs.

## Introduction

Plants perceive a wide range of glycan-derived structures through cell-surface receptors, using these molecular cues to monitor developmental processes, cell wall integrity, and interactions with microorganisms (Ngou et al. 2026). Glycans derived from plant and microbial extracellular matrices can therefore function as signals that inform plants about changes in their cellular environment. Recognition of these structures is mediated by distinct classes of plasma membrane-localized receptors, including lysin motif (LysM) receptor kinases, leucine-rich repeat malectin (LRR-MAL) receptor kinases and cysteine-rich receptor kinases (CRKs), which perceive structurally diverse glycan structures through their extracellular domain (Miya et al. 2007; Tseng et al. 2022; Martín-Dacal et al. 2023; Huang et al. 2026; Gysel et al. 2026). Depending on the nature and context of the recognized structure, glycan perception can result in developmental and cell-wall-associated responses, accommodation of symbiosis (van Boerdonk et al. 2024), or activation of plant immunity (Molina et al. 2024; Ngou et al. 2026). Here, we focus on the latter, namely glycan-derived signals that are perceived by plant immune receptors. In this context, glycan-derived molecules can function as microbe-associated molecular patterns (MAMPs) and damage-associated molecular patterns (DAMPs) that activate pattern-triggered immunity (PTI) through cell-surface pattern recognition receptors (PRRs; Ausubel 2005; Zipfel 2008; Yamaguchi and Huffaker 2011). PTI involves rapid responses such as cytosolic Ca²⁺ influx, mitogen-activated protein kinase (MAPK) activation, reactive oxygen species (ROS) production, and transcriptional reprogramming that collectively contribute to the establishment of cellular defense responses (Bigeard et al. 2015; Boutrot and Zipfel 2017). The cytosolic Ca²⁺ influx is one of the earliest PTI hallmarks and is widely used as a sensitive readout of immune activation by oligosaccharides (Ranf et al. 2011).

Plant and microbial extracellular polysaccharides can be degraded or remodeled during development and plant–microbe interactions, releasing short oligosaccharides, termed oligosaccharins (Albersheim and Darvill 1985; Darvill et al. 1992), that can be perceived by plant receptors. Examples include oligogalacturonides (OGs) derived from pectin, chitooligosaccharides derived from chitin or chitosan (deacetylated chitin), cello-oligosaccharides derived from cellulose, β-1,3-glucan oligosaccharides (Mélida et al. 2018), mixed-linkage glucan (MLG)-derived oligosaccharides, and oligosaccharides derived from structurally diverse hemicelluloses such as arabinoxylans, xyloglucans, xylans, and mannans (Sun et al. 2025).

These oligosaccharides differ in their composition, glycosidic linkages, degree of polymerization, and substitution patterns, features that can influence their recognition by plant receptors and their biological activity. However, how individual receptors selectively recognize structurally related oligosaccharides remains to be characterized.

Among the structurally diverse oligosaccharides perceived by plants, those derived from MLGs represent a particularly interesting class of β-glucans. MLGs are unbranched glucans composed of β-(1,4)-linked glucosyl stretches interrupted by β-(1,3)-linkages and are abundant components of the cell walls of grasses and other plants (Fincher 2009; Burton and Fincher 2009). MLGs have also been reported in bacteria, oomycetes and fungi, although their occurrence and distribution across microbial taxa remain to be fully elucidated (Fontaine et al. 2000; Pettolino et al. 2009; Pérez-Mendoza et al. 2015; Lampugnani et al. 2024; Menn et al. 2026). Structural heterogeneity of MLGs, including linkage pattern and distribution of β-(1,3)-linked residues, can give rise to structurally distinct oligosaccharides upon enzymatic degradation. Lichenase-mediated hydrolysis, for example, releases characteristic MLG-derived oligosaccharides that vary in degree of polymerization (DP) and linkage arrangement, such as G3G (MLG DP2; laminaribiose), G4G3G (MLG DP3; MLG43), and G4G4G3G (MLG DP4), where G denotes a glucosyl residue and the numbers indicate the linkage positions (Li et al. 2006; Chang et al. 2021). The resulting oligosaccharide profile depends on the enzyme: lichenases preferentially cleave β-(1,4) linkages adjacent to β-(1,3) linkages, whereas cellulases can generate products from β-(1,4)-linked regions (McCleary 1988; Rebaque et al. 2021; Kataoka et al. 2022). MLG-derived oligosaccharides such as G4G3G activate immune responses in *Arabidopsis thaliana* (*A. thaliana*) and other plant species, indicating that MLG-derived fragments can function as carbohydrate-derived immune signals (Barghahn et al. 2021; Rebaque et al. 2021; Martín-Dacal et al. 2023; Rebaque et al. 2025). This raises the question of how MLG-derived oligosaccharides are perceived by plant receptors.

Genetic studies in *A. thaliana* identified a group of *igp* (*impaired in glycan perception*) mutants with reduced immune responses to glycan-derived oligosaccharides. Among these, IGP1, also known as CORK1 (Cellulose Oligosaccharide Receptor Kinase 1; AT1G56145; Tseng et al. 2022), encodes a plasma membrane-localized LRR-MAL receptor kinase. Mutations in IGP1, as well as in the related IGP3 and IGP4 LRR-MAL receptor kinases, impair immune responses to selected MLG- and cellulose-derived oligosaccharides (Martín-Dacal et al. 2023), β-1,4-xylo-oligosaccharides, and XA^3^XX-arabinoxylan oligosaccharides (Fernández-Calvo et al. 2024), implicating these receptors in the perception of structurally diverse cell wall-derived oligosaccharides. Isothermal titration calorimetry (ITC) analyses have further shown that the purified *At*IGP1 extracellular domain (ECD) directly binds the cellulose-derived oligosaccharides cellotriose (CEL3) and cellopentaose (CEL5) with micromolar affinity, whereas binding to MLG DP3 (G4G3G) was not detected under the conditions tested (Martín-Dacal et al. 2023). This lack of detectable binding is notable because *igp1* mutants exhibit impaired immune responses to MLG-derived oligosaccharides, raising the possibility that other MLG-derived structures may be recognized by IGP1. Recent structural work has shown that purified *At*IGP1 ECD binds cello-oligosaccharides through a defined structural pocket in the LRR domain (Jiménez-Sandoval et al. 2026), but whether additional MLG-derived structures are recognized by IGP1 through this pocket or a different region has not been systematically evaluated.

Addressing this question requires a system in which receptor–oligosaccharide recognition can be monitored independently of downstream plant signaling and without receptor purification. Heterologous surface display, which presents receptor ECDs at the cell surface, offers such an approach. In yeast surface display (YSD), *Saccharomyces cerevisiae* (*S. cerevisiae*) is engineered to express the protein of interest as a fusion to Aga2p, which anchors it at the cell wall (Boder and Wittrup 1997). Epitope tags allow expression and surface localization of the fusion protein to be verified on intact cells, the cells can be used directly to test association with fluorescently labeled glycans, and specificity can be assessed by competition with unlabeled glycans. YSD has been successfully applied to characterize receptor-associated interactions and to engineer proteins with altered binding properties (Holler et al. 2000; Kariolis et al. 2014; Cherf and Cochran 2015). This approach therefore provides a complementary strategy for investigating the oligosaccharide recognition properties of the *At*IGP1 ECD directly at the cell surface.

In this study, we applied YSD to *At*IGP1 to investigate how structural features of MLG-derived oligosaccharides, particularly their degree of polymerization and glycosidic linkage arrangement, influence their association with the receptor ECD. By combining YSD-based interaction assays with measurements of cytosolic Ca²⁺ elevations following the application of these MLG-derived oligosaccharides to *A. thaliana* plants, we sought to assess their capacity to trigger immune responses and the dependence of these responses on IGP1. Together, these complementary approaches provide a framework for investigating the relationship between MLG-derived oligosaccharide association with *At*IGP1 and glycan-triggered immune activation in plants.

## Materials and Methods

### Cloning, yeast transformation and growth conditions

Two *A. thaliana* IGP1 (*At*IGP1; AT1G56145; PDB ID 9HHX) variants were cloned into the pYD1 Yeast Display Vector (Invitrogen) for heterologous expression and surface display in *S. cerevisiae*, resulting in Aga2p fusion proteins with an N-terminal Xpress epitope and a C-terminal V5 epitope and 6×His tag: IGP1+TM, comprising the ECD and the native transmembrane (TM) domain (amino acid residues 1-650), and IGP1-TM, containing the ECD only (amino acid residues 1-627).

A codon-optimized IGP1+TM DNA sequence was synthesized with flanking *Kpn*I and *Xho*I restriction sites by GeneArt (Thermo Fisher Scientific) and amplified by PCR using IGP1+TM_Fw1/IGP1+TM_Rv1 primers. IGP1-TM was amplified using the IGP1+TM template with IGP1+TM_Fw1/IGP1-TM_Rv1 primers. PCR products and the pYD1 vector were digested with *Kpn*I and *Xho*I (Thermo Fisher Scientific), ligated using T4 DNA ligase (New England Biolabs), and propagated in *E. coli* TOP10F′ cells under ampicillin selection. Constructs were verified by whole-plasmid sequencing (Microsynth Seqlab). Verified constructs were transformed into *S. cerevisiae* strain EBY100 using the lithium acetate method (Gietz and Schiestl 2007). Yeast cells were grown at 30 °C and 225 rpm using minimal dextrose (MD) medium supplemented with leucine and tryptophan (50 µg ml⁻¹ each) or without tryptophan for plasmid selection. Transformants were screened by colony PCR using pYD1_Fw1/IGP1_Rv1 and IGP1_Fw1/pYD1_Rv1, followed by Sanger sequencing. All primers used in this study are listed in Table S1.

### Yeast Surface Display

Surface display of *At*IGP1 fusion proteins was performed according to the pYD1 Yeast Display Vector Kit Manual (Invitrogen). Briefly, *S. cerevisiae* EBY100 transformants were cultivated in 0.67 % yeast nitrogen base supplemented with 0.5 % casamino acids (YNB-CAA) and 2 % glucose at 30 °C and 225 rpm to an OD₆₀₀ of 2–5. Cells were harvested by centrifugation at 3,000 × *g* for 5 min at 4 °C, washed twice with ddH₂O, resuspended in YNB-CAA supplemented with 2 % galactose at OD₆₀₀ of 0.5 and incubated for 48 h at 20 °C and 225 rpm.

### Western Blot Analysis

Two OD₆₀₀ units of *S. cerevisiae* cells were collected before (0 h) and after 48 h induction. Proteins were extracted in 1 × Laemmli buffer (Laemmli 1970) and denatured at 90 °C for 10 min. Protein extracts were separated on 8–16 % Mini-PROTEAN® TGX Stain-Free™ Precast Protein Gels (Bio-Rad) at 50 V for 10 min, followed by 200 V for 1 h. Total protein was visualized by UV activation in the ChemiDoc XRS+. Following, protein extracts were wet-transferred to PVDF membranes (Bio-Rad Cat. No. 162-0177) using a Mini Trans-Blot® Cell (Bio-Rad) at 100 V for 1 h at 4 °C. Membranes were blocked overnight at 4 °C in Tris-buffered saline (TBS) containing 2.4 % (w/v) bovine serum albumin (BSA), and incubated for 1 h at RT with mouse anti-V5 antibody (1:2,500; Thermo Fisher Scientific, Cat. No. R960-25) diluted in TBS-BSA supplemented with 0.1 % Tween-20 (TBST-BSA) with gentle shaking. After three washes with TBST, membranes were incubated for 1 h at RT with horseradish peroxidase (HRP)-conjugated goat anti-mouse IgG secondary antibody (1:25,000; Thermo Fisher Scientific, Cat. No. 31430). After two washes with TBST and one wash with TBS, immunoreactive signals were detected using the enhanced chemiluminescence (ECL) substrate (Thermo Fisher Scientific, Cat. No. 34577) and imaged with a ChemiDoc™ XRS+ system (Bio-Rad).

### Immunolabeling of *S. cerevisiae* cells

Surface-displayed *At*IGP1 was detected by immunolabeling of the C-terminal V5 epitope on intact *S. cerevisiae* cells. Two OD₆₀₀ units of induced cells were harvested by centrifugation at 3,000 × *g* for 5 min at 4 °C, washed twice with phosphate buffered saline (PBS; pH 6.5) and incubated for 1 h at 4 °C in PBS containing 1 mg ml⁻¹ BSA and FITC-conjugated anti-V5 monoclonal antibody (1:500; Thermo Fisher Scientific, Cat. No. 400-3) with vertical rotation at 9 rpm in the dark. Cells were washed three times with PBS and resuspended in 100 µl of PBS. 20 µl of the suspension were transferred onto a microscope slide for fluorescence image analysis.

### Oligosaccharides used in the experiments

MLG DP2 (G3G; laminaribiose; β-1,3-D-(Glc)₂; SMB01403-50MG), cellobiose (CEL2; β-1,4-D-(Glc)₂; C7252), and maltoheptaose (MAL7; α-1,4-D-(Glc)₇; M7753) were purchased from Sigma-Aldrich. Cellotriose (CEL3; β-1,4-D-(Glc)₃; #O-CTR), cellotetraose (CEL4; β-1,4-D-(Glc)₄; #O-CTE), cellopentaose (CEL5; β-1,4-D-(Glc)₅; #O-CPE), cellohexaose (CEL6; β-1,4-D-(Glc)₆; #O-CHE), MLG DP3 (G4G3G; β-1,4-D-(Glc)₂-β-1,3-D-Glc; #O-BGTRIB), MLG DP4 (G4G4G3G; β-1,4-D-(Glc)₃-β-1,3-D-Glc; #O-BGTETB), and hexaacetyl-chitohexaose (CHI6; β-1,4-D-(GlcNAc)₆; #O-CHI6) were purchased from Megazyme.

### Fluorescent labeling of oligosaccharides and oligosaccharide recognition analysis

Oligosaccharides were fluorescently labeled with Alexa Fluor 488 hydrazide (Thermo Fisher Scientific, Cat. No. A10436) by reductive amination using a twofold molar excess of dye relative to the oligosaccharide. Aniline in acetate buffer (pH 5.5) was added as a nucleophilic catalyst at a twofold molar excess relative to the dye and the reactions were incubated at 40 °C for 1 h in the dark. Sodium cyanoborohydride in DMSO was added at a 100-fold molar excess relative to the oligosaccharide, followed by incubation for an additional 1 h at 40 °C in the dark. All oligosaccharides were labeled in parallel under identical conditions. The reaction mixtures were diluted directly into the assay buffer. Concentrations of labeled oligosaccharide are therefore nominal, calculated from the amount of oligosaccharide used and assuming complete labeling, and the preparations contain residual free dye, reaction reagents and unlabeled oligosaccharide. Labeled oligosaccharides are denoted with the prefix “AF-“ (e.g., AF-CEL5). To control for dye- and reagent-associated background, AF-MAL7, labeled and handled identically, and untransformed wild-type (WT) cells were included in all assays.

Two OD₆₀₀ units of induced *S. cerevisiae* cultures were harvested by centrifugation at 3,000 × *g* for 5 min at 4 °C. Cell pellets were washed three times with PBS-YNB (1:1, v/v; PBS pH 6.5 and YNB medium) and subsequently resuspended in PBS-YNB containing 1.75 µM (nominal) fluorescently labeled oligosaccharide. Competition was assessed by incubating cells with fluorescently labeled oligosaccharide in the presence of a threefold molar excess of either the corresponding unlabeled oligosaccharide or MAL7 as a control. Samples were incubated at 4 °C for 16 h in the dark under vertical rotation at 9 rpm. Following incubation, cells were washed twice with PBS-YNB and resuspended in PBS-YNB for fluorescence microscopy.

### Fluorescence imaging and quantitative analysis of *S. cerevisiae* surface display and oligosaccharide recognition

Fluorescence and bright-field images were acquired using a Leica DM200 microscope equipped with a pE-300 white-light illumination system and an L5 filter cube (excitation 480/40 nm, emission 527/30 nm; Leica Microsystems). FITC and Alexa Fluor 488 fluorescence was detected with identical exposure settings for all samples. Fluorescence images were uniformly adjusted for brightness and contrast after acquisition for visualization purposes only. Quantification was performed on unadjusted images.

For each condition, 10 randomly selected fields of view were imaged per induction, from three independent biological inductions (30 fields in total). Cells were counted manually in the bright-field images, and a cell was scored as fluorescence-positive if its fluorescence exceeded a threshold defined from untransformed WT cells imaged with identical settings. Fluorescence-positive cells were expressed as a percentage of the total number of cells visible in the corresponding bright-field image. Surface display was quantified from anti-V5 fluorescence and oligosaccharide recognition from labeled oligosaccharide fluorescence. For both measurements, the mean percentage of fluorescent cells across the 30 WT fields imaged under the same condition was subtracted to correct for autofluorescence and nonspecific signals.

Because only a fraction of cells displayed the receptor, the percentage of oligosaccharide fluorescence-positive cells was normalized to the corresponding mean background-corrected surface display percentage of the respective variant (IGP1+TM: 17.8 %; IGP1-TM: 16.4 %), so that differences in display frequency between variants did not influence the comparisons.

Oligosaccharide Recognition (%) was calculated as:

(oligosaccharide-positive cells in the field [%] − mean oligosaccharide-positive cells in WT fields [%]) ÷ mean background-corrected surface display of the respective variant [%] × 100.

For example, a field of IGP1+TM cells with 4 AF-CEL5-positive cells out of 100 cells (4.0 %), a mean of 0.32 % in WT fields and a mean background-corrected surface display of 17.84 % yields a recognition of (4.0 − 0.32) ÷ 17.84 × 100 = 20.6 %.

Because this readout reports the proportion of positive cells and not the fluorescence intensity per cell, it is semi-quantitative.

### *Arabidopsis thaliana* growth conditions and detection of cytosolic Ca²⁺ responses by aequorin luminescence

Aequorin-expressing *A. thaliana* seedlings in the Col-0 background (Col-0*^AEQ^*; Knight et al. 1991; Ranf et al. 2011) and the *igp1-3^AEQ^* mutant background (Martín-Dacal et al. 2023) were grown for 8 d in liquid ½ Murashige and Skoog (MS) medium as previously described (Rebaque et al. 2021; Martín-Dacal et al. 2023). Cytosolic Ca²⁺ responses were monitored by aequorin luminescence using a Varioskan Lux Reader luminometer (Thermo Scientific) following treatment with CEL3 (100 µM), CEL5 (100 µM), MLG DP2 (G3G; 2.92 mM, corresponding to 1 mg ml⁻¹) or dH₂O (mock) as previously described for MLG-derived oligosaccharides (Rebaque et al. 2021; Martín-Dacal et al. 2023). Luminescence was recorded for 720 s after treatment. Total Ca²⁺ discharge was performed by treating seedlings with 2 M CaCl₂ in 20 % ethanol, and the Ca²⁺ burst was quantified as described previously (Martín-Dacal et al. 2023), and responses are expressed as a percentage of total Ca²⁺ discharge. Each treatment was analyzed in a total of 24 seedlings from 3 independent experiments.

### Statistical analysis

For quantification of surface display levels and oligosaccharide recognition individual measurements (fields of view; *n* = 30 per condition) from *N* = 3 independent inductions were pooled before statistical testing. Surface display levels (%) of IGP1+TM and IGP1-TM were compared by Welch’s *t*-test. Oligosaccharide recognition (%) was compared within each panel by one-way ANOVA followed by Tukey’s HSD post hoc test (*p* < 0.05); negative values resulting from background-correction were retained.

For Ca²⁺ responses of Col-0*^AEQ^* and *igp1-3^AEQ^*, individual measurement (seedlings; *n* = 24 per genotype and treatment) from *N* = 3 independent experiments were pooled before statistical testing. Total elicitor-induced cytosolic Ca^2+^ influx (Fig. 4E) of Col-0*^AEQ^* and *igp1-3^AEQ^* was compared for each treatment by unpaired two-tailed Student’s t-test.

## Results

### Engineered *S. cerevisiae* strains display *At*IGP1 protein variants on the cell surface

To establish a heterologous system for analysis of oligosaccharide recognition by *At*IGP1, we engineered *S. cerevisiae* to display the *At*IGP1 ECD on the yeast cell surface using an Aga2p-based system (Boder and Wittrup 1997). Two variants were generated to investigate whether the native TM domain affects ECD presentation: IGP1+TM, comprising the ECD and native TM domain, and IGP1-TM, comprising the ECD without native TM domain. Both variants were expressed as N-terminal Aga2p fusion proteins carrying a C-terminal V5 epitope tag for immunodetection of the displayed ECD (Fig. 1A).

**Figure 1:**
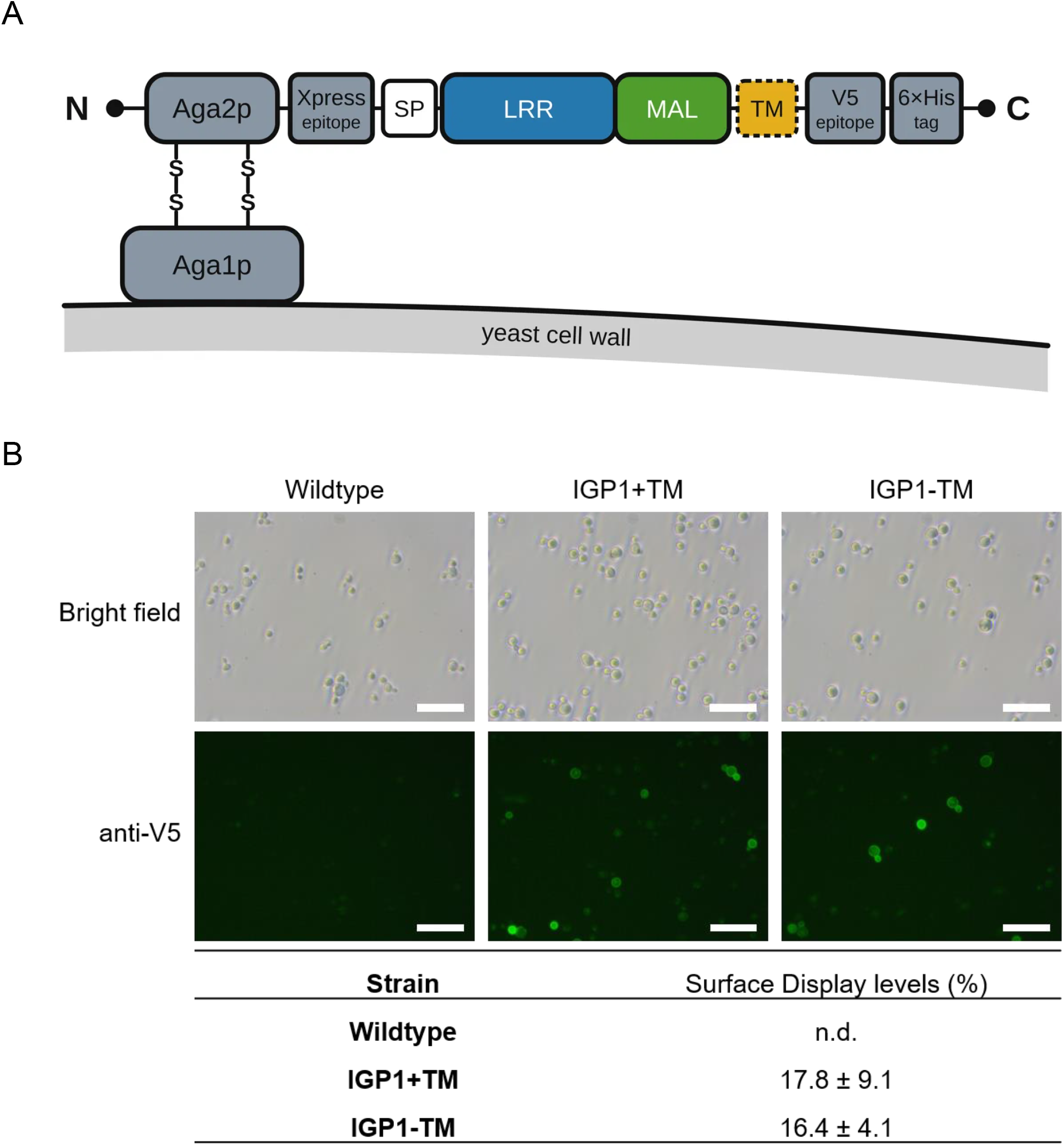
Surface display analysis of *At*IGP1 fusion proteins in *S. cerevisiae*. (A) Schematic representation (not to scale) of the *At*IGP1 fusion proteins surface-displayed on S. cerevisiae. *At*IGP1 is fused to Aga2p, which is linked to the cell wall-anchored Aga1p via disulfide bonds (S–S). The fusion proteins carry an N-terminal Xpress epitope and a C-terminal V5 epitope followed by a 6×His tag (N, N-terminus; C, C-terminus). IGP1+TM (residues 1–650) comprises the extracellular domain (ECD), which contains the leucine-rich repeat (LRR) and malectin (MAL) domains, and the native transmembrane domain (TM; dashed box); IGP1-TM (residues 1–627) comprises the ECD only. Residues are numbered from the native Met1 and include the native signal peptide (SP; residues 1–24). (B) Representative microscopy images of *S. cerevisiae* cells expressing *At*IGP1 fusion proteins with (IGP1+TM) or without (IGP1-TM) the native transmembrane domain, following immunolabeling using an FITC-conjugated anti-V5 monoclonal antibody. Non-transformed *S. cerevisiae* is shown as a control (Wildtype). Values in the table indicate the proportion of immunolabeled cells relative to the total population and are represented as mean ± SD. Quantification was performed using 30 fields of view per condition across three independent biological inductions *(N* = 3; *n* = 30). Values are background-corrected (mean of the untransformed wildtype fields subtracted from each field). Scale bars = 50 µm. n.d., not detected.

Following induction, surface localization of the fusion proteins was assessed by immunofluorescence labeling of the V5 epitope tag with a FITC-conjugated anti-V5 antibody on intact yeast cells. A fraction of cells expressing either IGP1+TM or IGP1-TM exhibited surface-associated fluorescence (Fig. 1B), indicating that the V5 epitope was accessible to the antibody on intact cells and supporting surface presentation of both *At*IGP1 variants. In contrast, induced untransformed WT cells displayed only background fluorescence, supporting the specificity of the immunolabeling.

As reported before for other YSD proteins (Yang et al. 2019; McKitrick et al. 2022), only a subset of the yeast population exhibited fluorescence above the defined background threshold. Nevertheless, surface display was consistently detected for both constructs across independent inductions. Quantification revealed that fluorescence-positive cells represented 17.8 % of the total cell population for IGP1+TM and 16.4 % for IGP1-TM (Fig. 1B). The proportion of fluorescence-positive cells did not differ significantly between the two variants (Welch’s t-test, *p* = 0.45), suggesting that the native TM domain did not substantially affect the proportion of cells displaying the *At*IGP1 ECD.

Expression of both IGP1 variants was further confirmed by SDS-PAGE and Western blot immunodetection of whole-cell extracts collected before (0 h) and after 48 h of induction (Fig. S1A, B). Anti-V5 immunoreactive bands were detected exclusively in induced transformants, whereas no signal was observed in untransformed WT cells or uninduced cultures. The fusion proteins migrated at apparent molecular masses around 100 kDa, exceeding their predicted sizes of ∼85 kDa for IGP1+TM and ∼83.5 kDa for IGP1-TM, which may reflect post-translational modification in yeast. IGP1-TM additionally showed a second immunoreactive protein at ∼85 kDa.

### Surface-displayed *At*IGP1 fusion proteins retain CEL5-specific oligosaccharide recognition

Having established surface presentation of the *At*IGP1 variants, we next investigated whether the displayed proteins retain specific oligosaccharide recognition previously described for the purified ECD. CEL5 was selected as a positive control because binding of the purified *At*IGP1 ECD to this cellulose-derived oligosaccharide has previously been demonstrated by ITC (Martín-Dacal et al. 2023).

Engineered cells displaying IGP1+TM or IGP1-TM were incubated in the presence of Alexa Fluor 488-labeled cellopentaose (AF-CEL5), and association was monitored by fluorescence microscopy (Fig. 2A). Fluorescently labeled maltoheptaose (AF-MAL7), a linear α-1,4-linked glucose heptamer, was included as a negative control (Fig. 2B). Both IGP1 variants exhibited pronounced fluorescence following incubation with AF-CEL5, whereas AF-MAL7 treatment resulted in fluorescence levels comparable to background. Untransformed WT control cells exhibited no detectable fluorescence with either labeled oligosaccharide, indicating negligible non-specific association of the fluorophore or oligosaccharides with the yeast cell surface.

**Figure 2:**
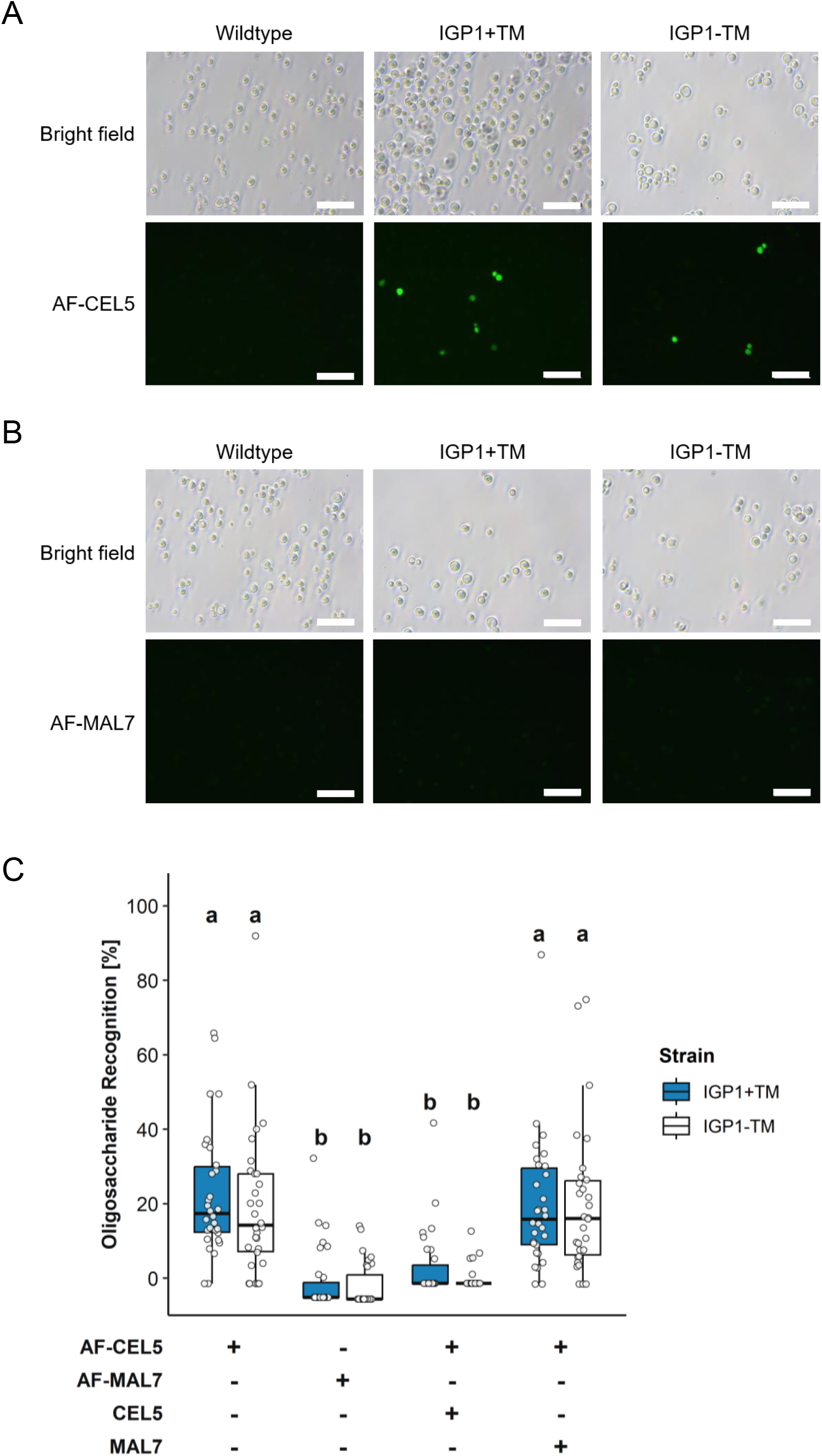
Oligosaccharide recognition of surface-displayed *At*IGP1 variants. (A, B) Representative fluorescence microscopy images showing recognition of Alexa Fluor 488-labeled cellopentaose (AF-CEL5; A) and Alexa Fluor 488-labeled maltoheptaose (AF-MAL7; B) at 1.75 µM (nominal) by yeast cells displaying *At*IGP1 variants (IGP1+TM and IGP1-TM). Brightness and contrast were adjusted uniformly for visualization purposes. Scale bars = 50 µm. C) Quantification of oligosaccharide recognition by *At*IGP1 variants in the presence of labeled and unlabeled competitor oligosaccharides (threefold molar excess). Incubation conditions are indicated below the x axis (+, present; -, absent). Box plots show the distribution of oligosaccharide recognition (%), corrected for background using untransformed WT cells and normalized to the surface display of the respective variant; center lines indicate medians, boxes represent the interquartile range (IQR), and whiskers extend to 1.5 × IQR. White circles represent individual measurements (*n* = 30 per condition across *N* = 3 independent biological replicates). Negative values result from background correction and were included in the statistical analysis. Different lowercase letters indicate statistically significant differences (one-way ANOVA with Tukey’s post hoc test, *p* < 0.05).

Background-corrected quantification of oligosaccharide recognition confirmed preferential recognition of AF-CEL5 over AF-MAL7 for both *At*IGP1 variants (Fig. 2C). The two *At*IGP1 variants exhibited comparable behavior, consistent with the surface-display analysis and indicating that removal of the native TM domain did not substantially alter CEL5 recognition in the YSD system.

To assess the specificity of the *At*IGP1–AF-CEL5 association, competition experiments were performed using unlabeled CEL5 or MAL7. Addition of a threefold molar excess of unlabeled CEL5 reduced AF-CEL5-associated fluorescence for both IGP1+TM- and IGP1-TM-expressing cells to background levels. In contrast, addition of an equivalent excess of unlabeled MAL7 did not significantly reduce the AF-CEL5-associated fluorescence. Thus, the fluorescent CEL5 signal could be competitively displaced by the corresponding unlabeled oligosaccharide but not by the control oligosaccharide MAL7 (Fig. 2C).

To define the oligosaccharide recognition specificity of the displayed *At*IGP1 variants, cells were incubated with additional fluorescently labeled oligosaccharides -namely the cello-oligosaccharides CEL2, CEL3, CEL4 and CEL6 and the chitin-derived hexaacetyl-chitohexaose (CHI6; Fig. S2B). Among these structures, only AF-CEL3 showed detectable association with both *At*IGP1 variants (Fig. S2A, B). Competition experiments were therefore also performed for CEL3: excess unlabeled CEL3 reduced AF-CEL3-associated fluorescence, whereas an equivalent excess of MAL7 reduced it partially but significantly for IGP1+TM, and did not reduce it significantly for IGP1-TM (Fig. S2C).

Together, these results show that surface-displayed *At*IGP1 selectively associates with CEL5 in a competitively displaceable manner and, less-specifically, with CEL3, while showing no detectable association with CEL2, CEL4, CEL6, CHI6, or MAL7 under the conditions tested.

### Surface-displayed *At*IGP1 shows specific association with MLG DP2

We next used the established yeast surface-display system to investigate recognition of mixed-linkage β-1,3/β-1,4-glucan (MLG)-derived oligosaccharides by the *At*IGP1 ECD. Based on the previously reported occurrence of DP2–DP4 structures among lichenase-released MLG oligosaccharides in the plant-associated fungus *Rhynchosporium secalis* (Pettolino et al. 2009), we selected fluorescently labeled MLG DP2 (G3G), DP3 (G4G3G) and DP4 (G4G4G3G), which differ in degree of polymerization and in the arrangement of their β-(1,3)- and β-(1,4)-linkages.

Yeast cells displaying IGP1+TM or IGP1-TM were incubated with fluorescently labeled MLG-derived oligosaccharides ranging from DP2 to DP4 (Fig. 3). Both *At*IGP1 variants exhibited detectable fluorescence following incubation with AF-MLG DP2, whereas incubation with longer MLG-derived oligosaccharides, DP3 and DP4, resulted in fluorescence levels comparable to background controls. Quantification of oligosaccharide recognition confirmed preferential association of both *At*IGP1 variants with MLG DP2 compared with DP3 and DP4.

**Figure 3:**
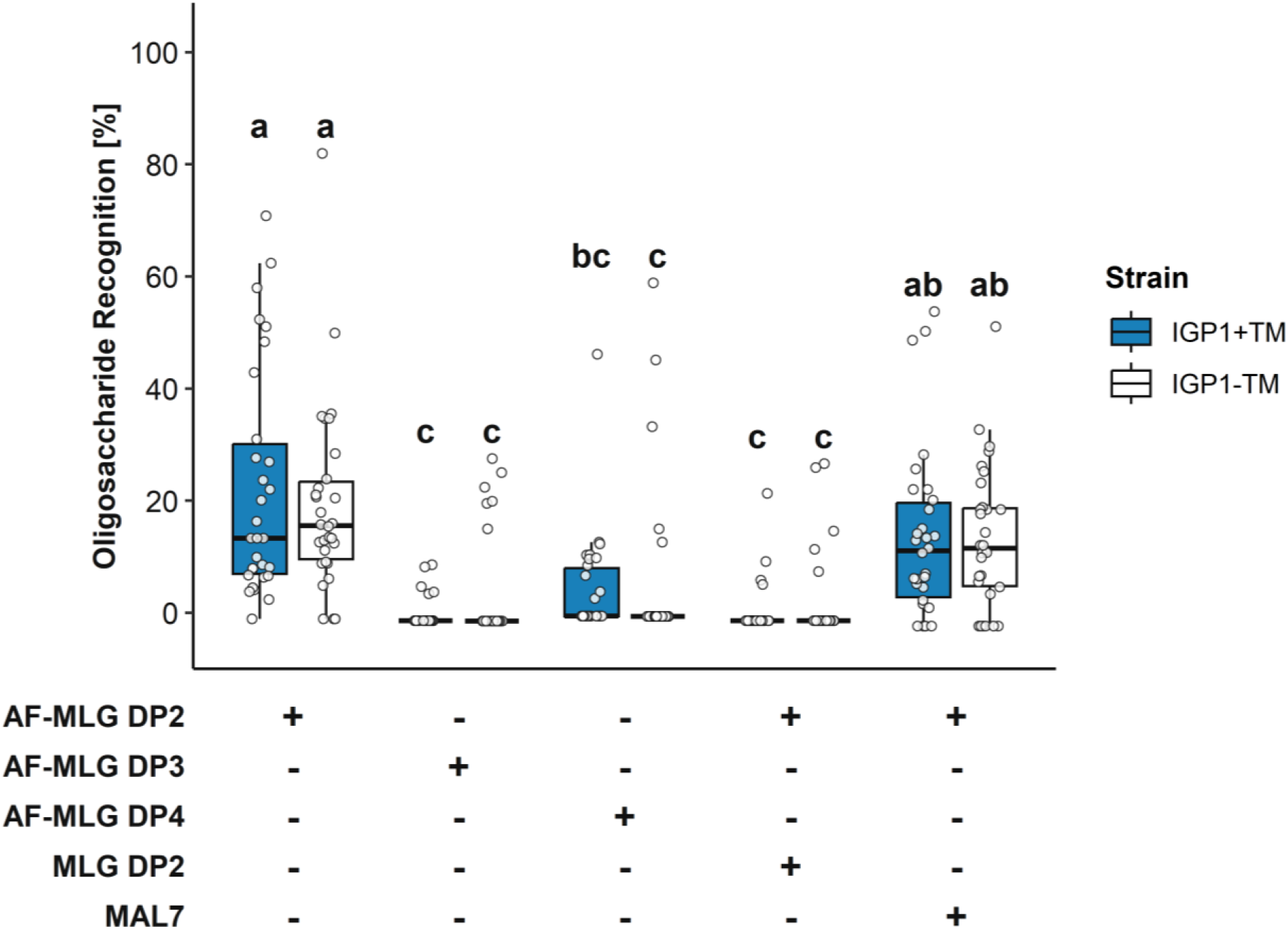
Selective recognition of MLG-derived oligosaccharides by surface-displayed *At*IGP1. Oligosaccharide recognition (%) of yeast surface-displayed *At*IGP1 variants in the presence of Alexa Fluor 488-labeled (AF-) MLG DP2 (G3G), DP3 (G4G3G) or DP4 (G4G4G3G) (1.75 µM, nominal) and of unlabeled competitor oligosaccharides (MLG DP2 or maltoheptaose, MAL7; threefold molar excess). Incubation conditions are indicated below the x axis (+, present; -, absent). Box plots show the distribution of oligosaccharide recognition (%), corrected for background using untransformed WT cells and normalized to the surface display of the respective variant; center lines indicate medians, boxes represent the interquartile range (IQR), and whiskers extend to 1.5 × IQR. White circles represent individual measurements (*n* = 30 per condition across *N* = 3 independent biological replicates). Negative values result from background correction and were included in the statistical analysis. Different lowercase letters indicate statistically significant differences (one-way ANOVA with Tukey’s post hoc test, *p* < 0.05).

To test whether the AF-MLG DP2 signal could be competitively displaced, we performed competition assays. For both variants, a threefold molar excess of unlabeled MLG DP2 significantly reduced the proportion of AF-MLG DP2-positive cells to levels comparable to those observed with AF-MLG DP3 and AF-MLG DP4, whereas an equivalent excess of MAL7 did not significantly alter the signal (Fig. 3). Notably, cellobiose (CEL2), which has the same degree of polymerization as MLG DP2 but a β-1,4 instead of a β-1,3 linkage, gave no detectable signal in the same assay (Fig. S2B), indicating that the association is not a general property of disaccharides but depends on the glycosidic linkage.

### Exogenous application of MLG DP2 induces an IGP1-dependent cytosolic Ca²⁺ response in ***Arabidopsis***

To test whether the association of MLG DP2 with the *At*IGP1 ECD observed in yeast is relevant *in planta*, we asked whether unlabeled MLG DP2 activates immune responses in *A. thaliana*. Because Ca²⁺ elevation is an early signaling event associated with activation of plant defense responses, we used the cytosolic Ca²⁺ response as a readout of immune activation. Ca²⁺ dynamics were monitored non-invasively in transgenic *A. thaliana* seedlings expressing the bioluminescent Ca²⁺ reporter aequorin in the Col-0 background (Col-0*^AEQ^*; Knight et al. 1991; Ranf et al. 2011). Treatment with MLG DP2 triggered a pronounced Ca²⁺ response in Col-0*^AEQ^* seedlings (Fig. 4C). In contrast to the rapid, single-peaked responses observed following CEL3 and CEL5 treatment (Fig. 4A, B), MLG DP2 elicited a delayed, biphasic cytosolic Ca²⁺ response, resembling similar kinetics as reported for MLG DP3 (G4G3G; Rebaque et al. 2021).

**Figure 4:**
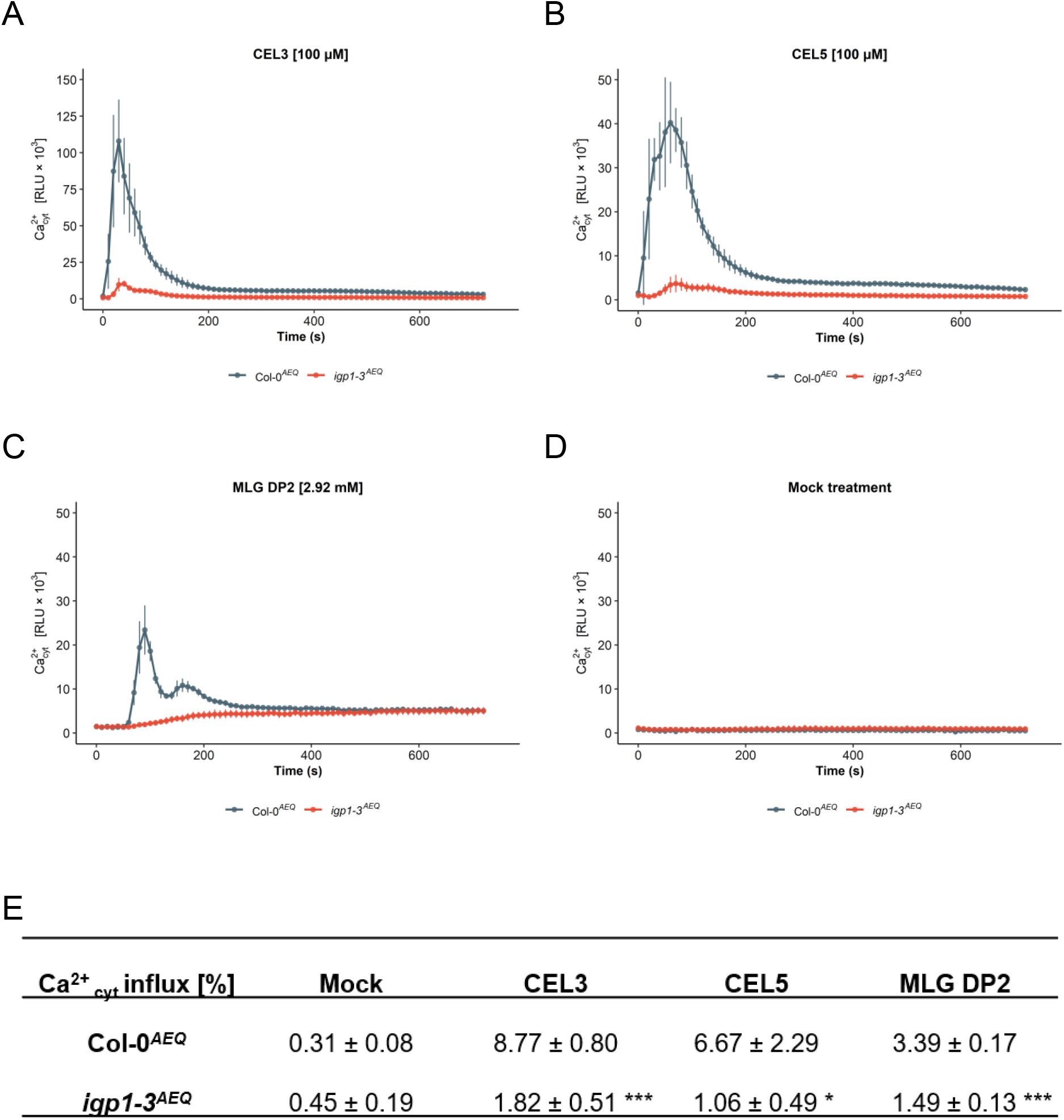
MLG DP2 elicits an immune-like cytosolic Ca²⁺ influx in *A. thaliana*. (A-D) Time-resolved cytosolic Ca²⁺ influx kinetics in Col-0*^AEQ^* and *igp1-3^AEQ^* reporter lines (AEQ, aequorin) following treatment with A) CEL3 (100 µM), B) CEL5 (100 µM), C) MLG DP2 (2.92 mM), or D) dH₂O (mock treatment). Cytosolic Ca²⁺ responses are displayed as relative luminescence units (RLU) over time. Data represent mean ± SEM (*n* = 24 seedlings per genotype and treatment from *N* = 3 independent experiments). Note the different y-axis scale in A. Total aequorin luminescence was discharged with CaCl₂ to determine percentage of elicitor-induced cytosolic Ca²⁺ influx. (E) Quantification of total elicitor-induced cytosolic Ca²⁺ influx for each treatment and genotype, presented as a percentage of total Ca²⁺-dependent luminescence obtained following CaCl₂ discharge. Data represent mean ± SEM (*n* = 24 seedlings per genotype and treatment). Asterisks indicate statistically significant differences between Col-0^AEQ^ and *igp1-3*^AEQ^ for each treatment individually, as determined by an unpaired two-tailed Student’s *t*-test (* = *p* < 0.05, *** = *p* < 0.001). SEM, standard error of the mean.

To assess the contribution of *At*IGP1, which was previously shown to be required for responses to cello- and MLG-derived oligosaccharides, the same treatments were performed in the *igp1-3^AEQ^* mutant (Martín-Dacal et al. 2023; Jiménez-Sandoval et al. 2026). Consistent with previous observations (Martín-Dacal et al. 2023; Jiménez-Sandoval et al. 2026), the Ca²⁺ transients triggered by CEL3 and CEL5 were markedly reduced in *igp1-3^AEQ^* (Fig. 4A, B). In *igp1-3^AEQ^* seedlings, the response to MLG DP2 was markedly reduced, with only a weak residual response remaining (Fig. 4C, D). Quantification of the normalized Ca²⁺ responses confirmed a strong reduction in MLG DP2-induced signaling in *igp1-3^AEQ^* compared with Col-0*^AEQ^* seedlings (Fig. 4E).

## Discussion

Our study establishes YSD as an approach for investigating oligosaccharide recognition by PRRs, using the *A. thaliana* receptor IGP1 as a proof of concept. In YSD, the receptor ECD is displayed on intact *S. cerevisiae* cells that can be incubated directly with labeled oligosaccharides and unlabeled competitors, which allows glycan recognition to be examined independently of the native plant signaling machinery and without receptor purification (Boder and Wittrup 1997, 2000; McKitrick et al. 2022; Lopez-Morales et al. 2023). Mutant and reporter assays *in planta*, in contrast, cannot distinguish receptor recognition from the contribution of additional receptors or downstream components. ITC provides quantitative affinities but requires the ECD to be purified in a properly folded state, which can be compromised by denaturation or aggregation during purification. In YSD, folding of the displayed ECD cannot be verified by such methods, but its oligosaccharide-recognition competence is supported by the association with CEL3 and CEL5 (see below). YSD thus complements *in planta* assays and ITC by testing receptor-glycan association in a heterologous context.

The platform was validated with the known *At*IGP1 ligands CEL3 and CEL5. Both surface-displayed *At*IGP1 variants, IGP1+TM and IGP1-TM, associated with the fluorescently labeled oligosaccharides, consistent with the binding of these oligosaccharides to the purified *At*IGP1 ECD (Martín-Dacal et al. 2023). Because ligand binding depends on the folded LRR–malectin pocket (Jiménez-Sandoval et al. 2026), this supports proper folding and functional presentation of the displayed ECD, which alone is sufficient for oligosaccharide recognition in the YSD platform. The association with labeled CEL5 was displaced by unlabeled CEL5 but not by MAL7, supporting its specificity. For CEL3, unlabeled CEL3 also reduced the signal, but MAL7 reduced it as well in IGP1+TM (Fig. S2C), indicating a less specific component; CEL5 therefore serves as the primary validation of the platform. In contrast, CEL4 and CEL6, which bind the purified ECD with affinities similar to CEL3 (Jiménez-Sandoval et al. 2026), showed no detectable association. Several factors may explain such missing signals. Rapidly dissociating interactions may be lost during washing, although the high density of displayed receptor on the cell surface may favor rebinding. The N-glycans of the ECD lie away from the binding pocket (Jiménez-Sandoval et al. 2026), but differences between plant and yeast glycosylation, with potential hypermannosylation of heterologous proteins in *S. cerevisiae* (Popova et al. 2023), and anchoring through Aga2p may still affect folding, orientation or ligand accessibility of the displayed ECD (Boder and Wittrup 1997). Fluorescent labeling through reductive amination, which opens the reducing-end glucosyl ring, may also reduce affinity, as oxidation of the reducing end abolishes CEL3 binding (Jiménez-Sandoval et al. 2026). A lack of signal thus does not exclude binding, whereas competitively displaced associations, as observed for CEL3 and CEL5, indicate glycan recognition. YSD is therefore best used as a semi-quantitative screening approach to identify candidate ligands for quantitative follow-up, for example by ITC with the purified ECD.

Among the further oligosaccharides tested, surface-displayed *At*IGP1 associated with the β-(1,3)-linked disaccharide MLG DP2 (G3G; laminaribiose), but not with CEL2, MLG DP3 (G4G3G), MLG DP4 (G4G4G3G), MAL7 or CHI6. The lack of association of MLG DP3 agrees with the absence of G4G3G binding to the purified ECD (Martín-Dacal et al. 2023). G3G was thus the only MLG-derived oligosaccharide tested that associated with *At*IGP1, which supports the possibility that MLG-derived structures other than G4G3G are recognized by the receptor.

Recent structural analyses revealed that the *At*IGP1 ECD binds CEL3 through a pocket containing three subsites accommodating β-(1,4)-linked glucosyl residues, consistent with the absence of detectable binding to cellobiose (Jiménez-Sandoval et al. 2026). Against this structural background, the observed association of fluorescently labeled G3G, a β-(1,3)-linked disaccharide, is unexpected. One possible explanation is that the Alexa Fluor 488 moiety attached to the reducing end contributes to this association, particularly given the accommodation of the reducing-end region within the binding pocket. However, the fluorophore alone is unlikely to account for the observed association, as similarly labeled CEL2, MAL7 and CHI6 showed no detectable association with the displayed receptor, whereas CEL3 and CEL5 did. Furthermore, competition with unlabeled G3G supports a contribution of the glycan itself to the observed interaction. Our data thus show an association of labeled G3G with the displayed *At*IGP1 ECD, but do not demonstrate that unmodified G3G binds the receptor with an affinity comparable to that of CEL3 or CEL5.

The potential recognition of G3G by *At*IGP1 raises questions about the biological relevance of this interaction. Although G3G is a structural motif found in MLG-derived oligosaccharides (Pettolino et al. 2009; Menn et al. 2026), it is not specific to MLG and may also be generated from other β-(1,3)-glucans, including plant callose (Ellinger and Voigt 2014) and fungal or oomycete glucans (Mélida et al. 2018). Thus, recognition of G3G would not necessarily represent the perception of an MLG-specific structural signature, but rather that of short β-(1,3)-glucan fragments of plant or microbial origin. Further, MLG is not detectable as a major cell wall component in *A. thaliana*, despite the ability of the plant to respond to MLG-derived oligosaccharides (Barghahn et al. 2021; Rebaque et al. 2021). Hence, MLG-derived structures encountered during plant–microbe interactions may originate from microbial cell walls, for example of *R. secalis* (Pettolino et al. 2009) or *Hyaloperonospora arabidopsidis* (Rebaque et al. 2021). Consistent with the perception of β-(1,3)-glucan oligosaccharides, *A. thaliana* responds to laminarihexaose (Mélida et al. 2018; Wanke et al. 2020), whereas long-chain laminarin does not trigger comparable responses (Wanke et al. 2020). If G3G is perceived through *At*IGP1, it should therefore activate immune responses in an IGP1-dependent manner.

Consistent with this prediction, application of unlabeled G3G to Col-0*^AEQ^* seedlings triggered a pronounced cytosolic Ca²⁺ elevation that was markedly reduced in *igp1-3^AEQ^*, indicating a substantial contribution of *At*IGP1. This extends previous genetic evidence that *igp1* mutants show impaired immune responses to MLG-derived oligosaccharides and could explain why these mutants are affected although the purified ECD does not bind G4G3G (Martín-Dacal et al. 2023). The residual response and the delayed, biphasic kinetics, which differed from the single-peaked responses to CEL3 and CEL5, are consistent with additional perception or signaling components, as reported for other β-glucans (Mélida et al. 2018; Wanke et al. 2020). Although an indirect contribution of *At*IGP1 cannot be excluded, given the millimolar concentration of G3G (2.92 mM; 1 mg ml⁻¹), about 30-fold above that of CEL3 and CEL5 (100 µM), the lack of dose-response analysis and of an inactive control structure, and the use of a single igp1 allele and an early readout, the association of G3G with the displayed ECD together with the substantially IGP1-dependent response *in planta* supports a contribution of *At*IGP1 to G3G perception.

In summary, we identified G3G (laminaribiose) as the only MLG-derived oligosaccharide of the tested panel that associates with the displayed *At*IGP1 ECD, and showed that it triggers a substantially IGP1-dependent Ca²⁺ response in *A. thaliana* seedlings. Although direct binding of unmodified G3G remains to be established through independent biochemical approaches such as ITC, our findings identify G3G as a candidate ligand of *At*IGP1. More broadly, our study demonstrates that YSD can be used to investigate receptor-glycan association separately from immune signaling, providing a straightforward entry point to link glycan-derived signals to the plant receptors that perceive them.

## Author contributions

VR and MP conceived and designed the project. BM performed experiments and analyzed data. BM, AM, LJ, VR and MP wrote the manuscript. All authors contributed to the article and approved the submitted version.

## Acknowledgments

This work was funded by the Cluster of Excellence on Plant Sciences (CEPLAS) by the Deutsche Forschungsgemeinschaft (DFG; German Research Foundation) under Germany’s Excellence Strategy – EXC 2048/1 – Project ID 390686111, and by grants PID2024-159175OB-I00 from MICIU/AEI/10.13039/501100011033 and by “ERDF/EU” (LJ, AM) and CEX2020-000999-S from Severo Ochoa Program for Centres of Excellence in R&D funded by MCIU/AEI/10.13039/501100011033 (AM).

## Data availability statement

All data supporting the findings in this study are available within the paper and its Supporting Information.

## Supporting information

**Figure S1.** Expression analysis of *At*IGP1 fusion proteins in *S. cerevisiae*.

**Figure S2.** Oligosaccharide recognition specificity of surface-displayed *At*IGP1 fusion proteins.

**Table S1.** Oligonucleotide primers used in this study.

## References

Albersheim, Peter, and Alan G. Darvill. 1985. “Oligosaccharins.” Scientific American 253 (3): 58–64.

Ausubel, Frederick M. 2005. “Are Innate Immune Signaling Pathways in Plants and Animals Conserved?” Nature Immunology 6 (10): 973–79.

Barghahn, Sina, Gregory Arnal, Namrata Jain, Elena Petutschnig, Harry Brumer, and Volker Lipka. 2021. “Mixed Linkage β-1,3/1,4-Glucan Oligosaccharides Induce Defense Responses in Hordeum Vulgare and Arabidopsis Thaliana.” Frontiers in Plant Science 12 (June): 682439.

Bigeard, Jean, Jean Colcombet, and Heribert Hirt. 2015. “Signaling Mechanisms in Pattern-Triggered Immunity (PTI).” Molecular Plant 8 (4): 521–39.

Boder, E. T., and K. D. Wittrup. 1997. “Yeast Surface Display for Screening Combinatorial Polypeptide Libraries.” Nature Biotechnology 15 (6): 553–57.

Boder, E. T., and K. D. Wittrup. 2000. “Yeast Surface Display for Directed Evolution of Protein Expression, Affinity, and Stability.” Methods in Enzymology 328: 430–44.

Boerdonk, Sarah van, Pia Saake, Alan Wanke, Ulla Neumann, and Alga Zuccaro. 2024. “β-Glucan-Binding Proteins Are Key Modulators of Immunity and Symbiosis in Mutualistic Plant-Microbe Interactions.” Current Opinion in Plant Biology 81 (102610): 102610.

Boutrot, Freddy, and Cyril Zipfel. 2017. “Function, Discovery, and Exploitation of Plant Pattern Recognition Receptors for Broad-Spectrum Disease Resistance.” Annual Review of Phytopathology 55 (1): 257–86.

Burton, Rachel A., and Geoffrey B. Fincher. 2009. “(1,3;1,4)-Beta-D-Glucans in Cell Walls of the Poaceae, Lower Plants, and Fungi: A Tale of Two Linkages.” Molecular Plant 2 (5): 873–82.

Chang, Shu-Chieh, Rebecka Karmakar Saldivar, Pi-Hui Liang, and Yves S. Y. Hsieh. 2021. “Structures, Biosynthesis, and Physiological Functions of (1,3;1,4)-β-D-Glucans.” Cells (Basel, Switzerland) 10 (3): 510.

Cherf, Gerald M., and Jennifer R. Cochran. 2015. “Applications of Yeast Surface Display for Protein Engineering.” Methods in Molecular Biology (Clifton, N.J.) (New York, NY), Methods in molecular biology (Clifton, N.J.), vol. 1319: 155–75.

Darvill, A., C. Augur, C. Bergmann, et al. 1992. “Oligosaccharins--Oligosaccharides That Regulate Growth, Development and Defence Responses in Plants.” Glycobiology 2 (3): 181–98.

Ellinger, Dorothea, and Christian A. Voigt. 2014. “Callose Biosynthesis in Arabidopsis with a Focus on Pathogen Response: What We Have Learned within the Last Decade.” Annals of Botany 114 (6): 1349–58.

Fernández-Calvo, Patricia, Gemma López, Marina Martín-Dacal, et al. 2024. “Leucine Rich Repeat-Malectin Receptor Kinases IGP1/CORK1, IGP3 and IGP4 Are Required for Arabidopsis Immune Responses Triggered by β-1,4-D-Xylo-Oligosaccharides from Plant Cell Walls.” Cell Surface (Amsterdam) 11 (100124): 100124.

Fincher, Geoffrey B. 2009. “Revolutionary Times in Our Understanding of Cell Wall Biosynthesis and Remodeling in the Grasses.” Plant Physiology 149 (1): 27–37.

Fontaine, T., C. Simenel, G. Dubreucq, et al. 2000. “Molecular Organization of the Alkali-Insoluble Fraction of Aspergillus Fumigatus Cell Wall.” The Journal of Biological Chemistry 275 (36): 27594–607.

Gietz, R. Daniel, and Robert H. Schiestl. 2007. “High-Efficiency Yeast Transformation Using the LiAc/SS Carrier DNA/PEG Method.” Nature Protocols 2 (1): 31–34.

Gysel, Kira, Simon Boje Hansen, Henriette Rübsam, et al. 2026. “Structural Basis for Size-Selective Perception of Chitin in Plants.” *Science (New York*, N.Y.) 394 (6821): 298–305.

Holler, P. D., P. O. Holman, E. V. Shusta, S. O’Herrin, K. D. Wittrup, and D. M. Kranz. 2000. “In Vitro Evolution of a T Cell Receptor with High Affinity for Peptide/MHC.” Proceedings of the National Academy of Sciences of the United States of America 97 (10): 5387–92.

Huang, Keyi, Jingni Wu, Yu Cao, et al. 2026. “A Pair of Chitosan Receptors Confer Broad-Spectrum Fungal Resistance.” Science (New York, N.Y.) 394 (6821): 291–97.

Jiménez-Sandoval, Pedro, Owen Kentish, Caroline Broyart, et al. 2026. “Glycan Recognition by a Plant Damage-Sensing Immune Receptor.” Science (New York, N.Y.), no. eaec6740 (September). 10.1126/science.aec6740.

Kariolis, Mihalis S., Yu Rebecca Miao, Douglas S. Jones 2nd, et al. 2014. “An Engineered Axl ‘decoy Receptor’ Effectively Silences the Gas6-Axl Signaling Axis.” Nature Chemical Biology 10 (11): 977–83.

Kataoka, Misumi, Han-Woo Kim, and Kazuhiko Ishikawa. 2022. “Recognition Mechanism of Endocellulase for β-Glucan Containing β(1 → 3),(1 → 4) Mixed-Linkages.” Carbohydrate Research 522 (108682): 108682.

Knight, M. R., A. K. Campbell, S. M. Smith, and A. J. Trewavas. 1991. “Transgenic Plant Aequorin Reports the Effects of Touch and Cold-Shock and Elicitors on Cytoplasmic Calcium.” Nature 352 (6335): 524–26.

Laemmli, U. K. 1970. “Cleavage of Structural Proteins during the Assembly of the Head of Bacteriophage T4.” Nature 227 (5259): 680–85.

Lampugnani, Edwin R., Kris Ford, Yin Ying Ho, et al. 2024. “Glycosyl Transferase GT2 Genes Mediate the Biosynthesis of an Unusual (1,3;1,4)-β-Glucan Exopolysaccharide in the Bacterium Sarcina Ventriculi.” Molecular Microbiology 121 (6): 1245–61.

Li, W., S. W. Cui, and Y. Kakuda. 2006. “Extraction, Fractionation, Structural and Physical Characterization of Wheat β-D-Glucans.” Carbohydrate Polymers 63 (3): 408–16.

Lopez-Morales, Joanan, Rosario Vanella, Elizabeth A. Appelt, et al. 2023. “Protein Engineering and High-Throughput Screening by Yeast Surface Display: Survey of Current Methods.” Small Science 3 (12): 2300095.

Martín-Dacal, Marina, Patricia Fernández-Calvo, Pedro Jiménez-Sandoval, et al. 2023. “Arabidopsis Immune Responses Triggered by Cellulose- and Mixed-Linked Glucan-Derived Oligosaccharides Require a Group of Leucine-Rich Repeat Malectin Receptor Kinases.” The Plant Journal: For Cell and Molecular Biology 113 (4): 833–50.

McCleary, Barry V. 1988. “Lichenase from Bacillus Subtilis.” In Methods in Enzymology. Elsevier.

McKitrick, Tanya R., Melinda S. Hanes, Charles S. Rosenberg, et al. 2022. “Identification of Glycan-Specific Variable Lymphocyte Receptors Using Yeast Surface Display and Glycan Microarrays.” Methods in Molecular Biology (Clifton, N.J.) (New York, NY), Methods in molecular biology (Clifton, N.J.), vol. 2421: 73–89.

Mélida, Hugo, Sara Sopeña-Torres, Laura Bacete, et al. 2018. “Non-Branched β-1,3-Glucan Oligosaccharides Trigger Immune Responses in Arabidopsis.” The Plant Journal: For Cell and Molecular Biology 93 (1): 34–49.

Menn, Benjamin, Klaus Pfeffer, Vicente Ramirez, and Markus Pauly. 2026. “Characterization of a Fungal Mixed-Linkage Glucan Synthase.” In bioRxiv. bioRxiv, September 18. 10.64898/2026.09.17.752320.

Miya, Ayako, Premkumar Albert, Tomonori Shinya, et al. 2007. “CERK1, a LysM Receptor Kinase, Is Essential for Chitin Elicitor Signaling in Arabidopsis.” Proceedings of the National Academy of Sciences of the United States of America 104 (49): 19613–18.

Molina, Antonio, Lucía Jordá, Miguel Ángel Torres, et al. 2024. “Plant Cell Wall-Mediated Disease Resistance: Current Understanding and Future Perspectives.” Molecular Plant 17 (5): 699–724.

Ngou, Bruno Pok Man, Yasuhiro Kadota, and Ken Shirasu. 2026. “Plant Cell Surface Receptors.” The Plant Journal: For Cell and Molecular Biology 125 (6): e70800.

Pérez-Mendoza, Daniel, Miguel Ángel Rodríguez-Carvajal, Lorena Romero-Jiménez, et al. 2015. “Novel Mixed-Linkage β-Glucan Activated by c-Di-GMP in Sinorhizobium Meliloti.” Proceedings of the National Academy of Sciences of the United States of America 112 (7): E757–65.

Pettolino, Filomena, Izumi Sasaki, Alisa Turbic, et al. 2009. “Hyphal Cell Walls from the Plant Pathogen Rhynchosporium Secalis Contain (1,3/1,6)-Beta-D-Glucans, Galacto- and Rhamnomannans, (1,3;1,4)-Beta-D-Glucans and Chitin.” The FEBS Journal 276 (14): 3698–709.

Popova, Larissa G., Dmitrii E. Khramov, Olga I. Nedelyaeva, and Vadim S. Volkov. 2023. “Yeast Heterologous Expression Systems for the Study of Plant Membrane Proteins.” International Journal of Molecular Sciences 24 (13): 10768.

Ranf, Stefanie, Lennart Eschen-Lippold, Pascal Pecher, Justin Lee, and Dierk Scheel. 2011. “Interplay between Calcium Signalling and Early Signalling Elements during Defence Responses to Microbe- or Damage-Associated Molecular Patterns.” The Plant Journal: For Cell and Molecular Biology 68 (1): 100–113.

Rebaque, Diego, Cristian Carrasco-López, Parvathy Krishnan, et al. 2025. “Zymoseptoria Tritici Stealth Infection Is Facilitated by Stage-Specific Downregulation of a β-Glucanase.” The New Phytologist 248 (6): 3191–207.

Rebaque, Diego, Irene Del Hierro, Gemma López, et al. 2021. “Cell Wall-Derived Mixed-Linked β-1,3/1,4-Glucans Trigger Immune Responses and Disease Resistance in Plants.” The Plant Journal: For Cell and Molecular Biology 106 (3): 601–15.

Sun, Guangzheng, Yu Xiao, Heng Yin, Ke Yu, Yiming Wang, and Yuanchao Wang. 2025. “Oligosaccharide Elicitors in Plant Immunity: Molecular Mechanisms and Disease Resistance Strategies.” Plant Communications 6 (12): 101469.

Tseng, Yu-Heng, Sandra S. Scholz, Judith Fliegmann, et al. 2022. “CORK1, A LRR-Malectin Receptor Kinase, Is Required for Cellooligomer-Induced Responses in Arabidopsis Thaliana.” Cells (Basel, Switzerland) 11 (19): 2960.

Wanke, Alan, Hanna Rovenich, Florian Schwanke, et al. 2020. “Plant Species-Specific Recognition of Long and Short β-1,3-Linked Glucans Is Mediated by Different Receptor Systems.” The Plant Journal: For Cell and Molecular Biology 102 (6): 1142–56.

Yamaguchi, Yube, and Alisa Huffaker. 2011. “Endogenous Peptide Elicitors in Higher Plants.” Current Opinion in Plant Biology 14 (4): 351–57.

Yang, Xiaoyu, Hongting Tang, Meihui Song, Yu Shen, Jin Hou, and Xiaoming Bao. 2019. “Development of Novel Surface Display Platforms for Anchoring Heterologous Proteins in Saccharomyces Cerevisiae.” Microbial Cell Factories 18 (1): 85.

Zipfel, Cyril. 2008. “Pattern-Recognition Receptors in Plant Innate Immunity.” Current Opinion in Immunology 20 (1): 10–16.

