## Supplementary Figures for "Yeast surface display reveals structure-dependent β-glucan association with the plant immune receptor *At*IGP1"

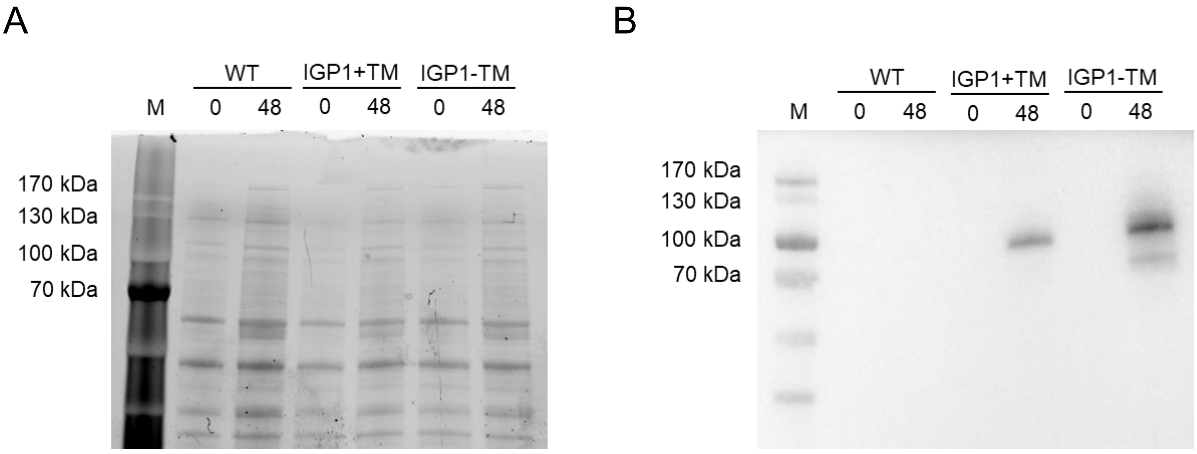


**Supplementary Figure S1: Expression analysis of *At*IGP1 fusion proteins in *S. cerevisiae*.**
(A) SDS-PAGE analysis showing total protein profiles from non-induced (0 h) and induced (48 h) yeast strains expressing IGP1+TM or IGP1-TM, alongside untransformed wild-type (WT) cells, visualized via UV transillumination.
(B) Western blot analysis of corresponding cell extracts detecting *At*IGP1 fusion proteins using an anti-V5 antibody. Molecular weight markers (M; in kDa) are indicated on the left in both panels. Lanes are labeled with the time after induction (h).

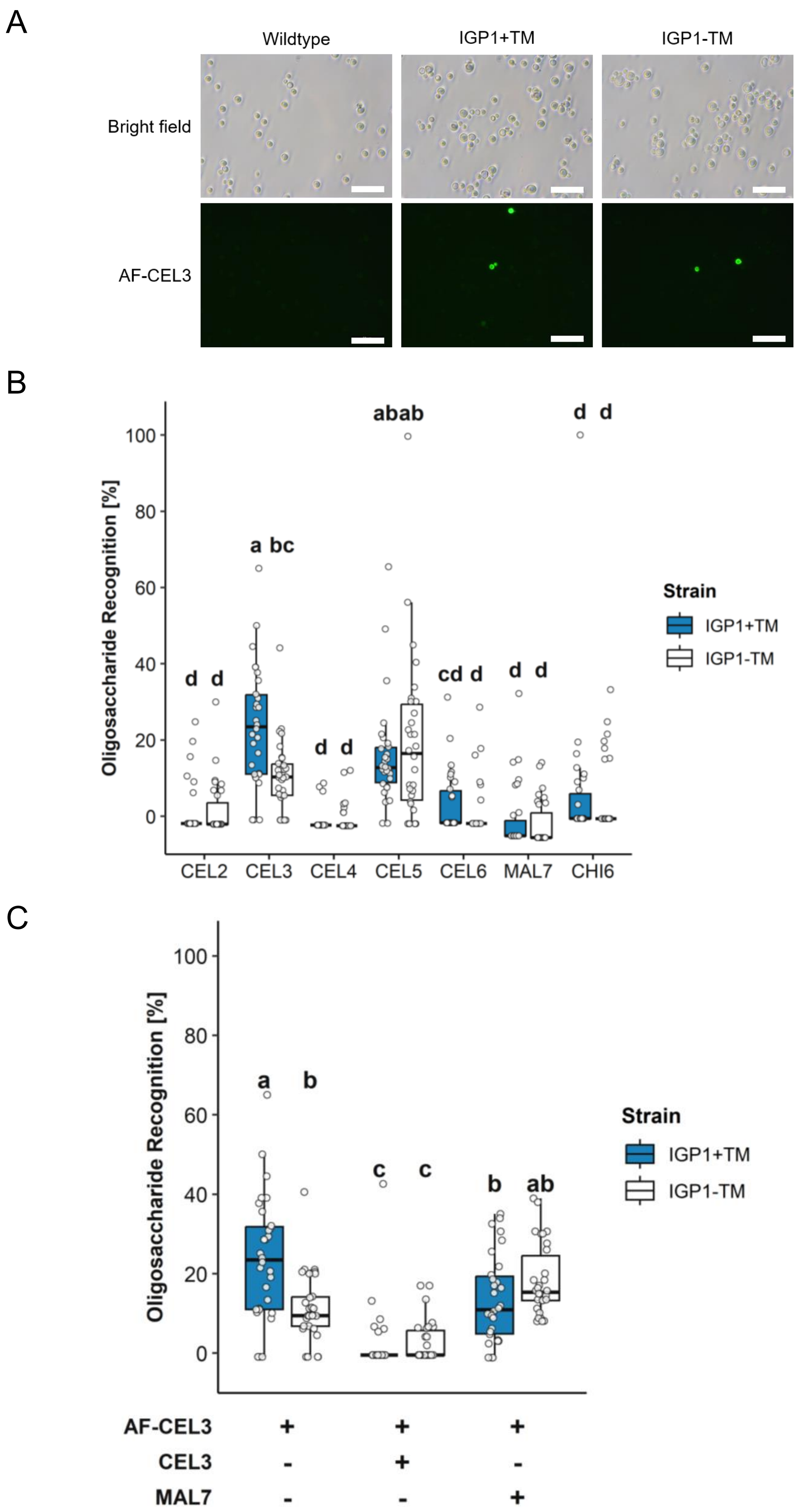


**Supplementary Figure S2: Oligosaccharide recognition specificity of surface-displayed *At*IGP1 fusion proteins.**

(A) Representative fluorescence microscopy images showing recognition of Alexa Fluor 488-labeled cellotriose (AF-CEL3; 1.75 µM (nominal)) by yeast displaying *At*IGP1 variants (IGP1+TM and IGP1-TM). Scale bars = 50 µm.

(B,C) Oligosaccharide recognition (%) of yeast surface-displayed *At*IGP1 variants. B) Recognition of fluorescently labeled oligosaccharides. C) Competition assay using fluorescently labeled cellotriose (AF-CEL3; 1.75 µM, nominal) and unlabeled competitor oligosaccharides (threefold molar excess). Incubation conditions are indicated below the x axis (+, present; -, absent). Box plots show the distribution of oligosaccharide recognition (%), corrected for background using untransformed WT cells and normalized to the surface display of the respective variant; center lines indicate medians, boxes represent the interquartile range (IQR), and whiskers extend to 1.5 × IQR. White circles represent individual measurements ($n$ = 30 per condition across $N$ = 3 independent biological replicates). Negative values result from background correction and were included in the statistical analysis. Different lowercase letters indicate statistically significant differences (one-way ANOVA with Tukey's post hoc test, $p < 0.05$).
