## Supplementary Tables for "Yeast surface display reveals structure-dependent β-glucan association with the plant immune receptor *At*IGP1"

### Supplementary Material:

#### Supplementary Table S1. Oligonucleotide primers used in this study

| Primer name | Sequence (5’ to 3’) | Purpose | Template/Target |
|---|---|---|---|
| **pYD1_Fw1** | GCTGCTAGTACGGATTAGAAG | Colony PCR and sequencing of transformed constructs | pYD1+IGP1+TM; pYD1+IGP1-TM |
| **pYD1_Rv1** | GTCGATTTTGTTACATCTACAC | Colony PCR and sequencing of transformed constructs | pYD1+IGP1+TM; pYD1+IGP1-TM |
| **IGP1_Fw1** | GTCAGGTGAAATTCCAAGCTC | Colony PCR and sequencing of transformed constructs | pYD1+IGP1+TM; pYD1+IGP1-TM |
| **IGP1_Rv1** | CATGACGGAAGGTCACCTG | Colony PCR and sequencing of transformed constructs | pYD1+IGP1+TM; pYD1+IGP1-TM |
| **IGP1+TM_Fw1** | ATCTGTACGACGATGACGATAAGGT | Amplification of synthesized IGP1+TM construct | Synthetic fragment IGP1+TM |
| **IGP1+TM_Rv1** | CTAGACTCGAGCAACAGGATAG | Amplification of synthesized IGP1+TM construct | Synthetic fragment IGP1+TM |
| **IGP1-TM_Rv1** | CTAGACTCGAGGTTCTTCTTACTCTT | Creation of IGP1-TM amplicon for restriction digestion cloning | Synthetic fragment IGP1+TM |
